# Modeling collaterally sensitive drug cycles: shaping heterogeneity to allow adaptive therapy

**DOI:** 10.1101/2020.07.02.184952

**Authors:** Nara Yoon, Nikhil Krishnan, Jacob Scott

**Affiliations:** Adelphi University; Case Western Reserve University; Cleveland Clinic

## Abstract

In previous work, we focused on the optimal therapeutic strategy with a pair of drugs which are collaterally sensitive to each other, that is, a situation in which evolution of resistance to one drug induces sensitivity to the other, and vice versa. [1] Here, we have extended this exploration to the optimal strategy with a collaterally sensitive drug sequence of an arbitrary length, *N*(≥ 2). To explore this, we have developed a dynamical model of sequential drug therapies with *N* drugs. In this model, tumor cells are classified as one of *N* subpopulations represented as {*R_i_*|*i* = 1,2,…, *N*}. Each subpopulation, *R_i_*, is resistant to ‘*Drug i*’ and each subpopulation, *R*_*i*–1_ (or *R_N_*, if *i* = 1), is sensitive to it, so that R_i_ increases under ‘*Drug i*’ as it is resistant to it, and after drug-switching, decreases under ‘*Drug i* + 1’ as it is sensitive to that drug(s).

Similar to our previous work examining optimal therapy with two drugs, we found that there is an initial period of time in which the tumor is ‘shaped’ into a specific makeup of each subpopulation, at which time all the drugs are equally effective 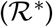. After this shaping period, all the drugs are quickly switched with duration relative to their efficacy in order to maintain each subpopulation, consistent with the ideas underlying adaptive therapy. [2, 3]

Additionally, we have developed methodologies to administer the optimal regimen under clinical or experimental situations in which no drug parameters and limited information of trackable populations data (all the subpopulations or only total population) are known. The therapy simulation based on these methodologies showed consistency with the theoretical effect of optimal therapy.

## 1 Introduction

Despite the development of a large pharmacopoeia of novel anti-cancer drugs, curative treatments remain elusive after systematic, or metastatic, spread of cancer. The evolution of resistance to initially effective therapies is one of the primary forces behind this phenomenon. This evolution is a complex phenomenon influenced by a variety of factors and their interactions [4, 5, 6], including genetic mutation and changed frequency of gene expression [7, 8, 9], drug efflux pumps on the cell membrane [10, 11], tumor microenvironement [12] and so on. Despite the difficulty of elucidating these complicated mechanisms, a multitude of (epi)genetic factors may converge to evolve finite phenotypes. Resistance to a particular drug can represent each of these phenotypes. Furthermore, such diversity in resistance phenotype can be leveraged to find synergistic combinations in which resistance and sensitivity factors of the involved drugs are properly engaged, like multiple cogwheels rolling together. Drugs having such relationships are called collaterally sensitive drugs or negatively cross resistant drugs. Particularly, there is utility in a drug sequence which completes a cycle of such relationships. (e.g., the three drugs connected by the red arrows in Figure 1.) With such cycles, one could, in theory, generate infinitely long drug sequences which can be used in long term therapy to mitigate the evolution of resistance in a tumor. [13] In light of this, we extend our previous work on two-drug cycles [1] to arbitrary length cycles in this paper.

**Figure 1:**
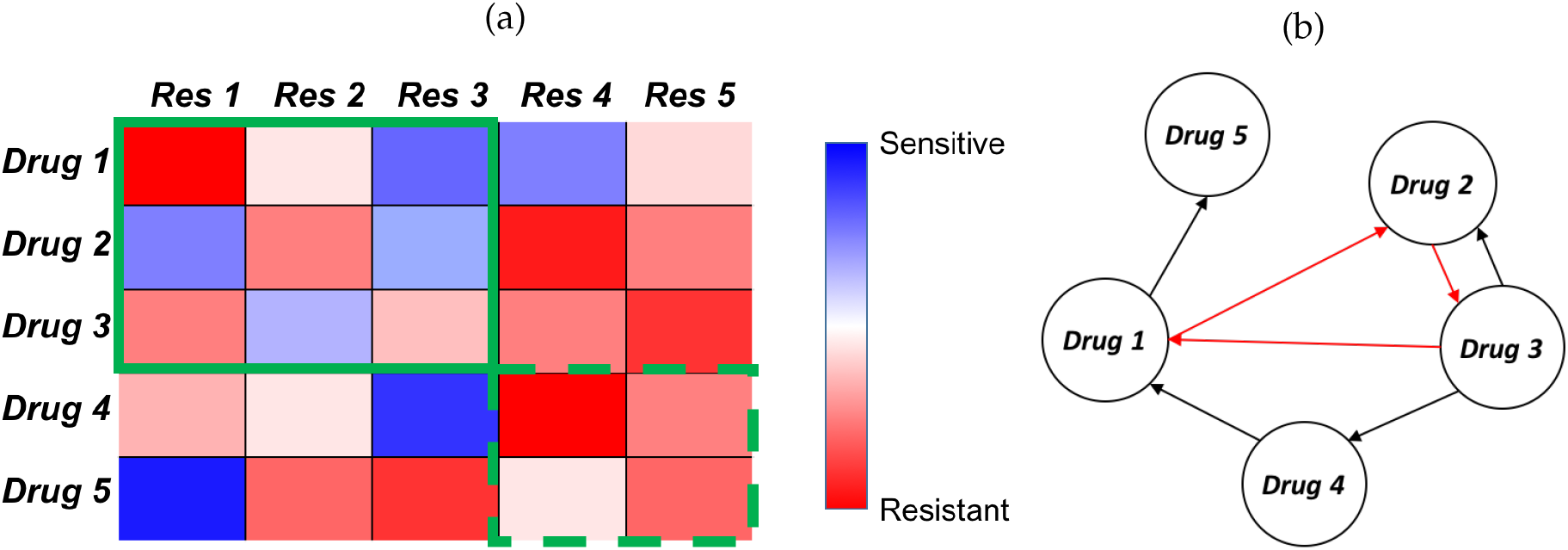
A hypothetical collateral sensitivity map (CSM), and associated collateral sensitivity network (CSN) among 5 drugs. (a) A collateral sensitivity map: This map represents the results of a hypothetical experiment in which tumors are exposed to one drug (row), and after resistance to this drug develops are tested against another drug (column). The color is then the *change* in sensitivity from the wild type to the evolved strain. (b) A collateral sensitivity network showing every drug pair and their collateral sensitivity relationship. Nodes represent drugs, and directed edges point from drugs, which when resistance develops, end with sensitivity to another (edge terminus). The four drugs connected by the red arrows is an example of collaterally sensitive drug cycle.

Tumors consists of diverse cells in terms of cellular traits (heterogeneity in genotypes and phenotypes) and/or surrounding environment (affinity to blood vessels, fibroblasts, etc.) which has been comprehensively reviewed. [14] This heterogeneity has been captured in numerous ways mathematically using a variety of different formalisms [15]. One type of simple model germane to the work here represents dimensionless fractions of finite number of cell subpopulations, assuming their total population is fixed. Tracking of the frequency of subpopulations has been efficiently used to explore the dynamics of tumors and therapy in the setting of evolutionary game theory [16, 17, 18, 19]. Other models described the population dynamics of also finitely many cell types [20, 21, 22], that can simulate the dynamical behavior of both subpopulations and total populations of cancers.

Some other modeling work has incorporated more detailed heterogeneous framework, by introducing continuously changing cellular biology [23, 24] or stromal microenvironment environment [25, 26]. In this paper, we chose to account for the intermediate level of modeling complexity. Using constrained ODEs, as in EGT, but also considering differential drug effect, similar to the concept of fitness landscapes [27]. With that, we defined a drug efficacy measure based on cell population, and explored the effects of collaterally sensitive drug schedules.

For parsimony and analytical tractability, we assume that there are possibly three fitness levels under each drug: lowest (for most sensitive cells), highest (for most resistant cells) and intermediate (for other neutral cells). Mutations in this population structure result only in progressively higher fitness values, specifically from sensitive to neutral and neutral to resistant types. We make no fine grained biological assumptions for the purposes of this work, but assume only that a fitness metric of overall proliferation ability in the process of convergent evolution [28, 29, 30, 31]. A subpopulation having a same fitness value is assumed to be homogeneous in terms of how it responds to drug exposure. And, while it has been shown that fitness can often change as a function of drug dose, so called seascapes [32], and that this can be useful for control, we do not consider that possibility here [33].

In this study we ask the following questions: Given knowledge of the evolutionary patterns of resistance, what is the optimal method of using a large panel of drugs? What can we learn about the evolution of resistance from observing patient outcomes? Can each patient be their own control?

The remainder of this manuscript is structured as follows. The details of our model are described in Section 2. Based on the model, we derive the optimal treatment strategy and a practical method of its implementation, which are discussed in Section 3 and Section 4 respectively. Our finding of optimal treatment is consistent with the concept of ‘minimum effective dosage’ in the adaptive therapy paradigm, which optimizes drug effectiveness with the least risk of drug resistance development [2]. Section 5 includes a discussion of adaptive therapy, as well as overall conclusions and discussions of this work.

## 2 Modeling for collaterally sensitive drug cycles

Based on our previous model of collateral sensitivity cycles in two drugs [1], we have developed an extended model for an arbitrary length of *N* drugs cycles, *Drug* 1, *Drug* 2, …, and *Drug N*. In the model, tumor cells are classified into *N* subpopulations, *R*_1_, *R*_2_,…, and *R_N_*. Each subpopulation, *R_i_* is resistant under *Drug i*, sensitive under *Drug i*′ (see Table 1 for the definition of *i*′), and neutral under any of other drugs. Therefore, we can simulate the patterns of collateral sensitivity sequences in terms of the resistant cell populations (for example, *R_i_*′) under a particular drug (i.e., *Drug i*′ in the example) which subsequently declines under the next drug in the cycle to which it is sensitive (i.e., *Drug i*).

**Table 1:** Definitions of parameters and variables related to System (1)

| variables/parameters | descriptions |
| --- | --- |
| $N$ | number of drugs in cycle |
| $i$ | index of a chosen drug, $i \in \{1, 2, \dots, N\}$ |
| $i'$ | index of the drug whose resistant factor is sensitive to *Drug* $i$,<br>$i - 1$ for $i \geq 2$, and $N$ for $i = 1$ |
| $t$ | time variable |
| $[0, t_{shape}]$ | time interval of shaping period |
| $R_i$ | cell population resistant under *Drug* $i$, sensitive under *Drug* $i + 1$<br>(for $i < N$) or *Drug* 1 (for $i = N$) and neutral under all the other drugs |
| $\mathcal{R}$ | vector of subpopulations, $\mathcal{R} = (R_1 \ \dots \ R_N)^T$ |
| $\mathcal{R}^*$ | population fractions (i.e., $\sum_{k=1}^N R_k^* = 1$) at which all the drugs<br>are equally effective, $\mathcal{R}^* = (R_1^* \ \dots \ R_N^*)^T$ |
| $TP(t)$ | total population, $TP(t) = \sum_{k=1}^N R_k(t)$ |
| $TPD(t)$ | derivative of total population, $TPD(t) = TP'(t)$ |
| $p_r^i$ | proliferation rate of cells resistant to *Drug* $i$ |
| $p_s^i$ | proliferation rate of cells sensitive to *Drug* $i$ |
| $p_0^i$ | proliferation rate of cells neutral to *Drug* $i$ |
| $\mathcal{P}^i \in \mathbb{R}^N$ | vector of proliferation rates under *Drug* $i$,<br>$\mathcal{P}^i := (p_j^i) \text{ with } p_j^i = \begin{cases} p_r^i & \text{if } j = i \\ p_s^i & \text{if } j = i' \\ p_0^i & \text{otherwise} \end{cases}$ |
| $g_s^i \ (\overline{g_s^i})$ | transition rate from sensitive to all neutral types (each neutral type)<br>under *Drug* $i$, $\overline{g_s^i} := g_s^i / (N - 2)$ |
| $g_0^i$ | transition rate from neutral to resistant types under *Drug* $i$ |
| $\{\lambda_r^i, \lambda_s^i, \lambda_0^i\}$ | per capita turnover rates for the three types of cells under *Drug* $i$,<br>$\lambda_r^i := p_r^i, \lambda_s^i := p_s^i - g_s^i, \lambda_0^i := p_0^i - g_0^i$ |
| $\mathcal{D}(i) \in \mathbb{R}^{N \times N}$ | matrix of rate parameters in population dynamics under *Drug* $i$,<br>$\mathcal{D}(i) := (d_{j,k}^i) \text{ with } d_{j,k}^i = \begin{cases} \lambda_r^i & \text{if } j = k = i \\ \lambda_s^i & \text{if } j = k = i' \\ \lambda_0^i & \text{if } j = k \notin \{i, i'\} \\ \overline{g_0^i} & \text{if } j = i \text{ and } k \notin \{i, i'\} \\ \overline{g_s^i} & \text{if } k = i' \text{ and } j \notin \{i, i'\} \\ 0 & \text{otherwise} \end{cases}$ |
| $ef_i$ | effect of *Drug* $i$ |
| $\widetilde{ef}_i$ | approximated effect of *Drug* $i$ |

Under each drug (*Drug i*), we assigned three types of total proliferation rates (“birth rate”-“death rate”), for resistant 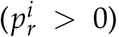, sensitive 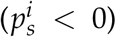 and neutral 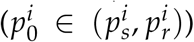) cells. Assuming evolution of each cellular type occurs toward higher fitness levels (i.e., proliferation rates in our model), we accounted for two kinds of transitions: from sensitive to neutral types 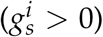 and neutral to resistant types 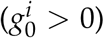. The dynamics of *R*_1_, *R*_2_, …, and *R_N_* under any chosen drug, *Drug i*, are described in Figure 2.

**Figure 2:**
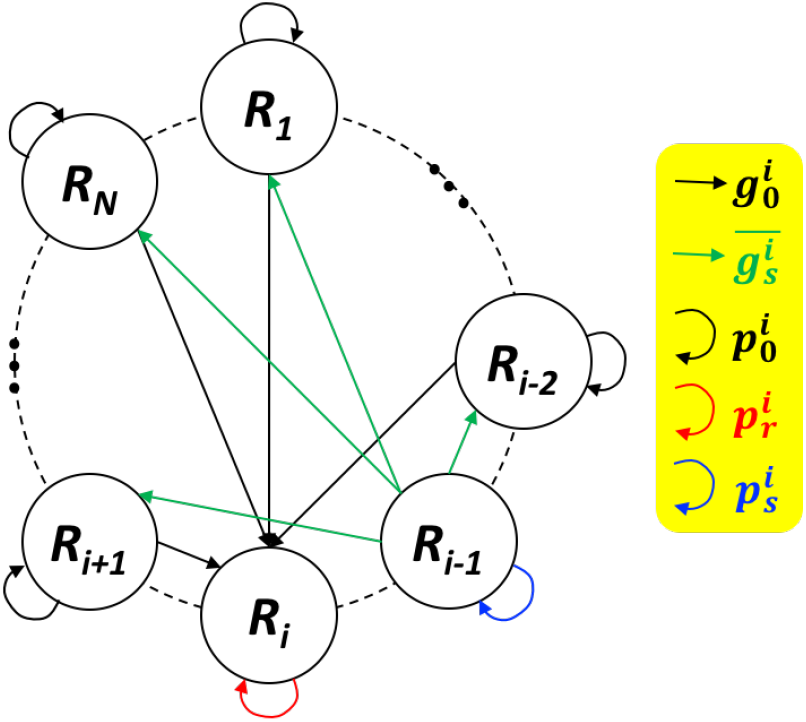
Population dynamics under *Drug i* **therapy**, where *R_i_* and *R*_*i*–1_ (*i* – 1 = *i*′ ≥ 1) are resistant and sensitive cell populations respectively, with all the other compartments being populations of neutral type. The left panel shows the schematic of transitions/turnovers among *R_j_*s, and the right panel shows the associated system of differential equations. Here, 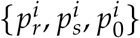 are proliferation rate of resistant, sensitive, and neutral cells, 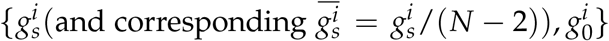 are transition rates from sensitive to neutral and, from neutral to resistant types.

In our study, system (1) is used to describe the dynamics of cell populations under a single drug. To study the dynamics under multiple drugs switched over time, we switch the

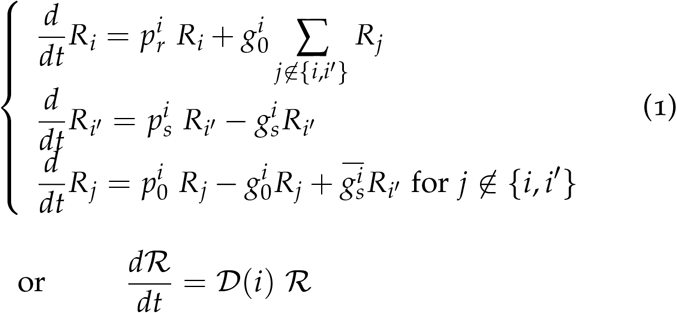

drug index, *i*, in the system accordingly. The resulting piece-wise continuous differential system will describe the effect of this drug switch strategy. An example of population histories with 4 collaterally sensitive drugs switched as indicated is shown in Figure 3.

**Figure 3:**
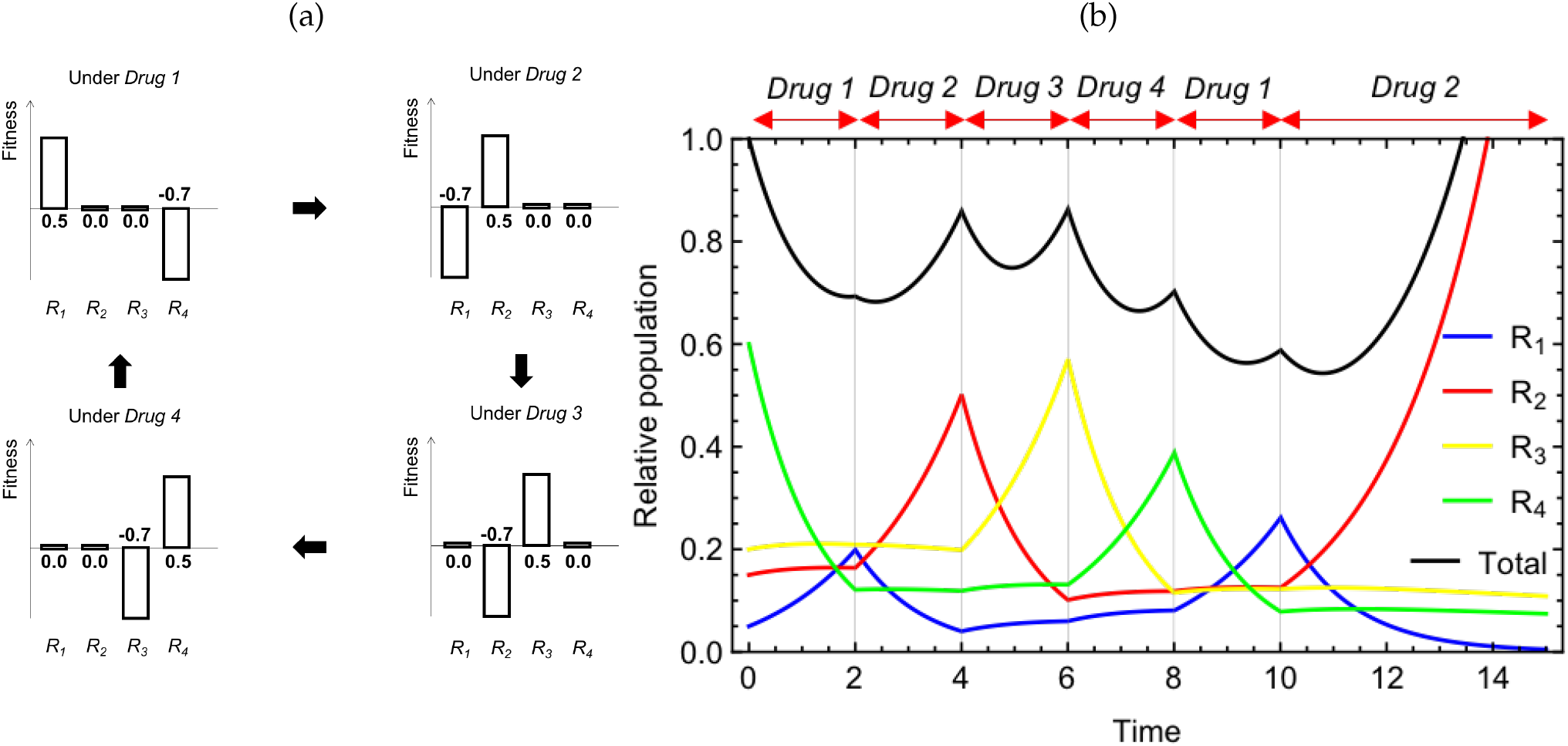
Example simulation with hypothetical drugs that complete a collateral sensitivity cycle. (a) Demonstration on the fitness levels of the cell populations classified in terms of sensitivity and resistance under the collaterally sensitive drugs. (b) Simulated effect of a sequence of the drugs based on the cell classification and fitness levels from (a) (i.e., {*p_r_*, *p_s_*, *p*_0_} = {0.5, −0.7,0.0} for all drugs). Other used values are {*g_s_*, *g*_0_} = {0.1,0.05} (transition rates) for all drug and 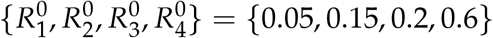 (initial populations).

Rapid drug rotation with chosen intensities of the *N* drugs (i.e., *f_i_*Δ*t*-long with *Drug i* where 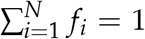 and Δ*t* → 0^+^) is employed in our optimal therapeutic strategy. This will be described in detail in the next section. In our modeling framework, the fast switch is highly related to the therapy with a drug mixture, since its corresponding cell population model is in a similar form as the single drug dynamics (Equation (1)). The only difference is the transition matrix which is replaced by the linear combination of the matrices of all the drugs 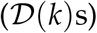 with the relative intensities (*f_k_*s), as described below.

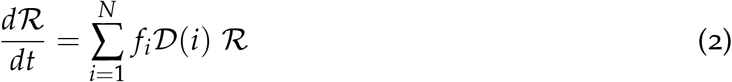

For the derivation of the Equation (2), see Theorem A.6 in Appendix A.

## 3 Optimal therapeutic schedule

### 3.1 Description of the optimal population dynamics

Analysis of a system comprised by multiple systems of (1) with different values of *i*, *p^i^*s and *g^i^*s, is limited due to the complexity of the system. Instead, we studied it numerically to find the optimal therapeutic strategy. To define control, we chose to minimize the area under the total population curve over a chosen time interval [0, *x*] for some time *x* > 0,

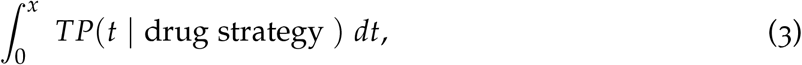

for varying drug administrations. Our numerical study determined the treatment regimen for the best possible effects to minimize (3). It can be achieved when the best drug(s) is chosen at every given moment (*t* = *t*_1_). Here, the best drug(s) at *t*_1_ (e.g., *Drug i*) means the drug(s) which have the highest effectiveness in decreasing total population (such that *ef_i_*(*t*_1_) ≤ *ef_i_*(*t*_1_) for any *j* ∈ {1,2,…, *N*}), where the formulation of the drug effect is defined in following way

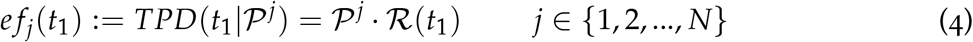

for *Drug j* at *t*_1_. 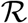 under the optimal strategy obeys the following system,

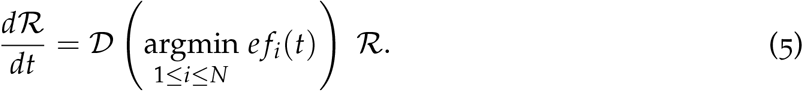

While we are deriving a mathematically optimal therapy schedule, we realize it is impractical to instantaneously switch drugs in realistic clinical situations. Therefore, we developed a simulation algorithm to choose and accordingly switch to an effective drug at a given discrete time step 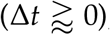, which is described in Figure 4 (a) – where a longer Δ*t* could be considered for specific clinical situations. An example of the optimal administration simulated with the discrete scheme, compared to a choice of non-optimal administration which changes drugs at minimum time points of total population, is shown in Figure 4 (b, c). The panel (b) shows tumor size, which is smaller with the optimal therapy over the time than the size with the non-optimal therapy, and the panel (c) shows how much the optimal therapy is better, measured by (3), in this example.

**Figure 4:**
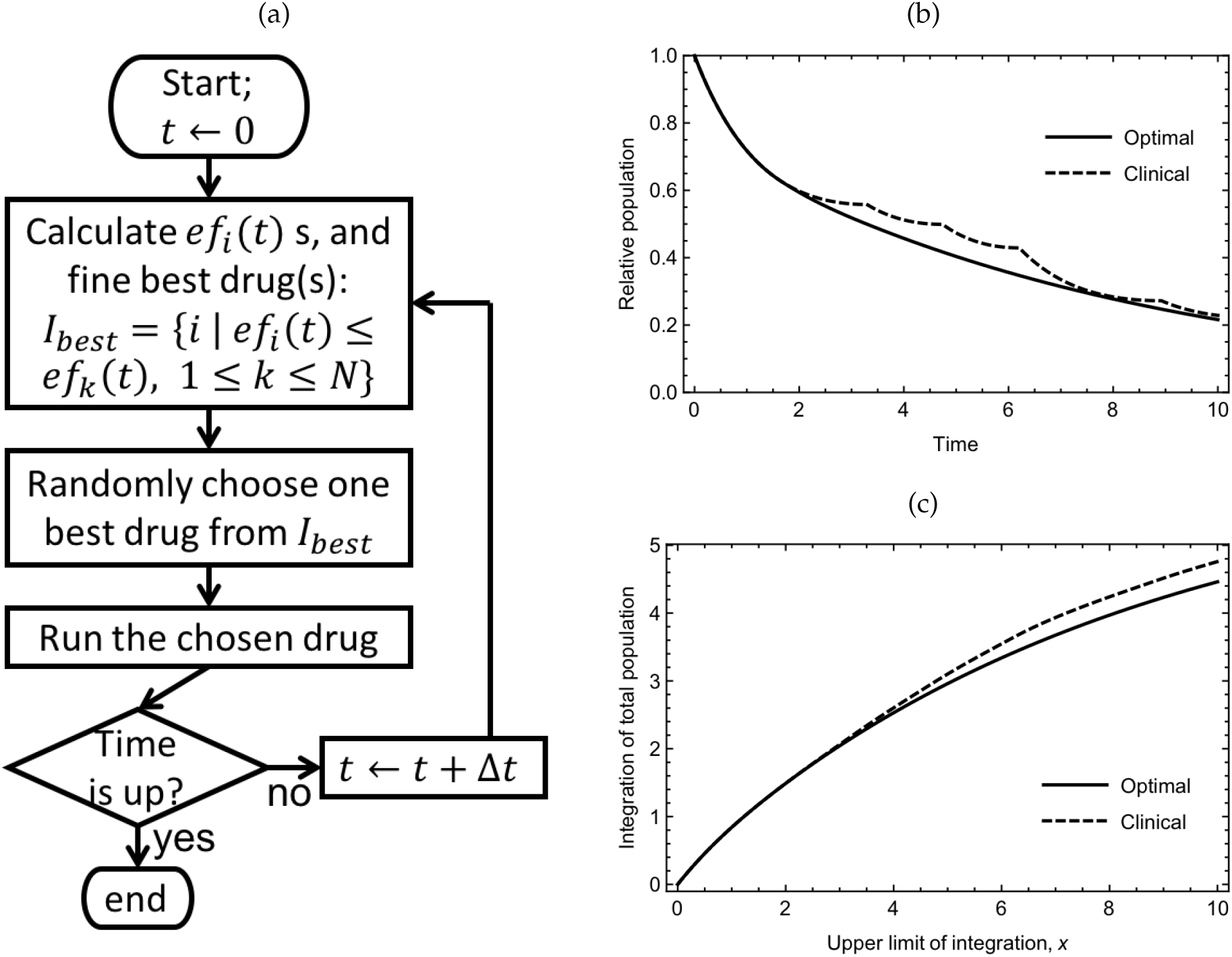
Algorithm for practical realization of the optimal therapeutic strategy (System (5)). (a) Flow chart of the optimal therapy algorithm. (b, c) Dynamical trajectories of the optimal strategy (solid curves) simulated through (a), compared to the trajectories of an example strategy relevant to the best possible standard clinical approach (minimum-point switches; dashed curves). The total populations of time histories is shown at Panel (b), and the integrations of total populations (3) from *t* = 0 to varying upper limit (x-axis) is shown in Panel (c). Parameters/conditions are: {*p_r_*, *p_s_*, *p*_0_} = {0.2,0.7,0.0} and {*g_s_*,*g*_0_} = {0.05,0.05} for all drugs, and 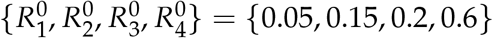.

### 3.2 Properties of the optimal therapy

In this section, we will study several properties about the optimal therapeutic regimen based on two examples of “symmetric" drug cycle (Figure 5) and an “asymmetric" drug cycle (Figure 6). We define symmetric drug cycle as a set of drugs which have an identical parameter value for each type of dynamical event, i.e., 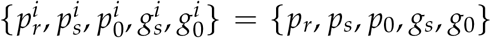 for all *i*, and an asymmetric drug cycle as having different parameter values for at least one type of dynamical event, i.e., 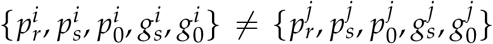 for some *i ≠ j*. We will begin with the simpler case of symmetric drug cycles first and then generalize our observations to asymmetric drug cycles.

**Figure 5:**
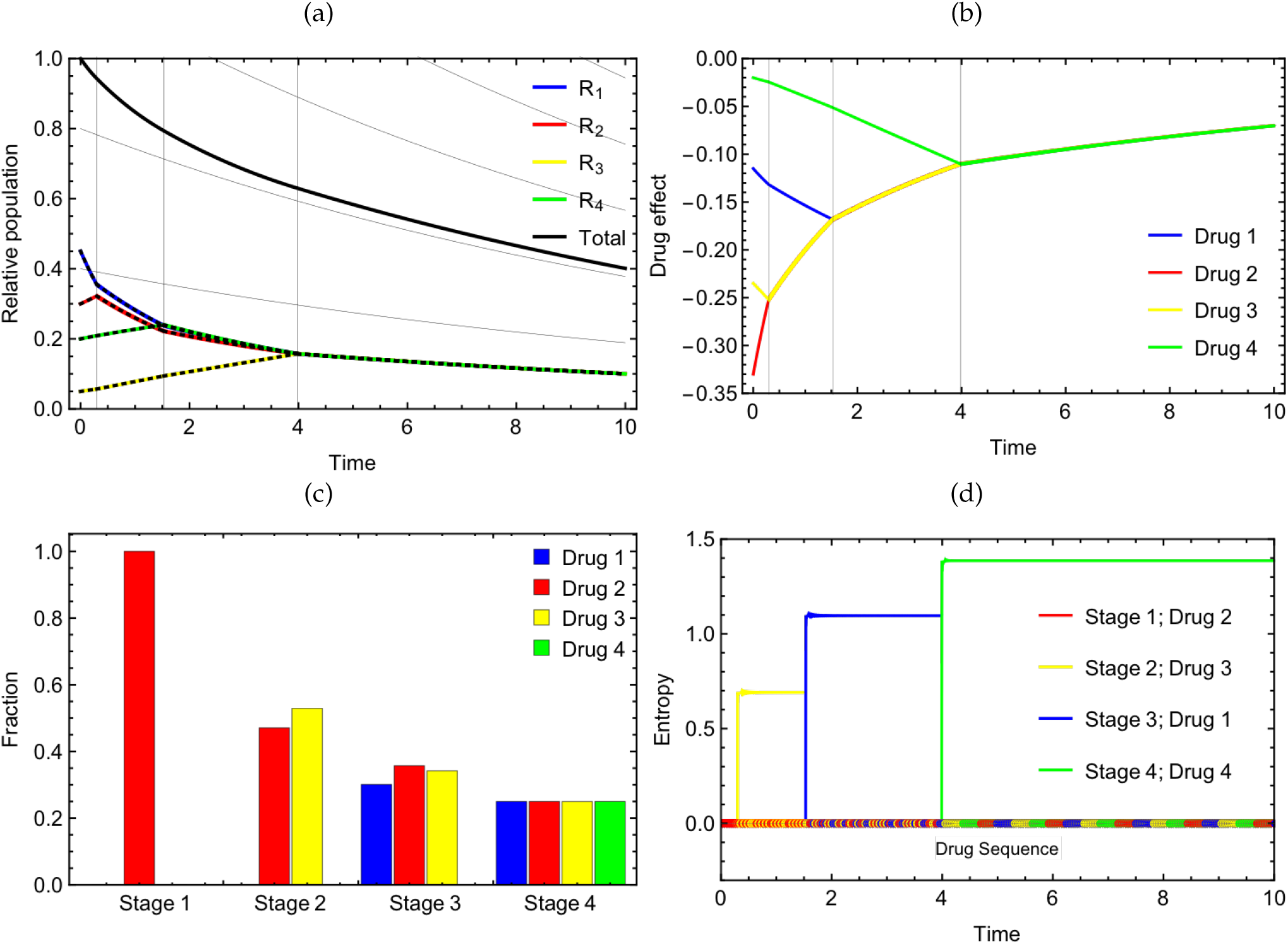
Optimal therapeutic dynamics with a cycle of four symmetric drugs. (a) Histories of subpopulations and total population with gray lines/curves indicating the ends of stages and exponential curves compatible to the last stage. (b) Effects of the drugs (rate of total population change under the drugs) measured by (4) along the optimal population dynamics. (c) Stage-wise relative frequencies of the drugs chosen in the optimal histories simulation by the algorithm (Figure 4 (a)). (d) Shannon entropy change over sequences of chosen drug indices starting from the beginning of each stage to the time points (x-axis). Selected drugs at every time points are indicated by the colored dots along the line of *y* = 0. Parameters for all panels are {*p_r_, p_s_, p*_0_} = {0.2,0.7,0.1}, {*g_s_,g*_0_} = {0.1,0.05}, 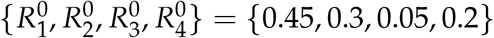

**Figure 6:**
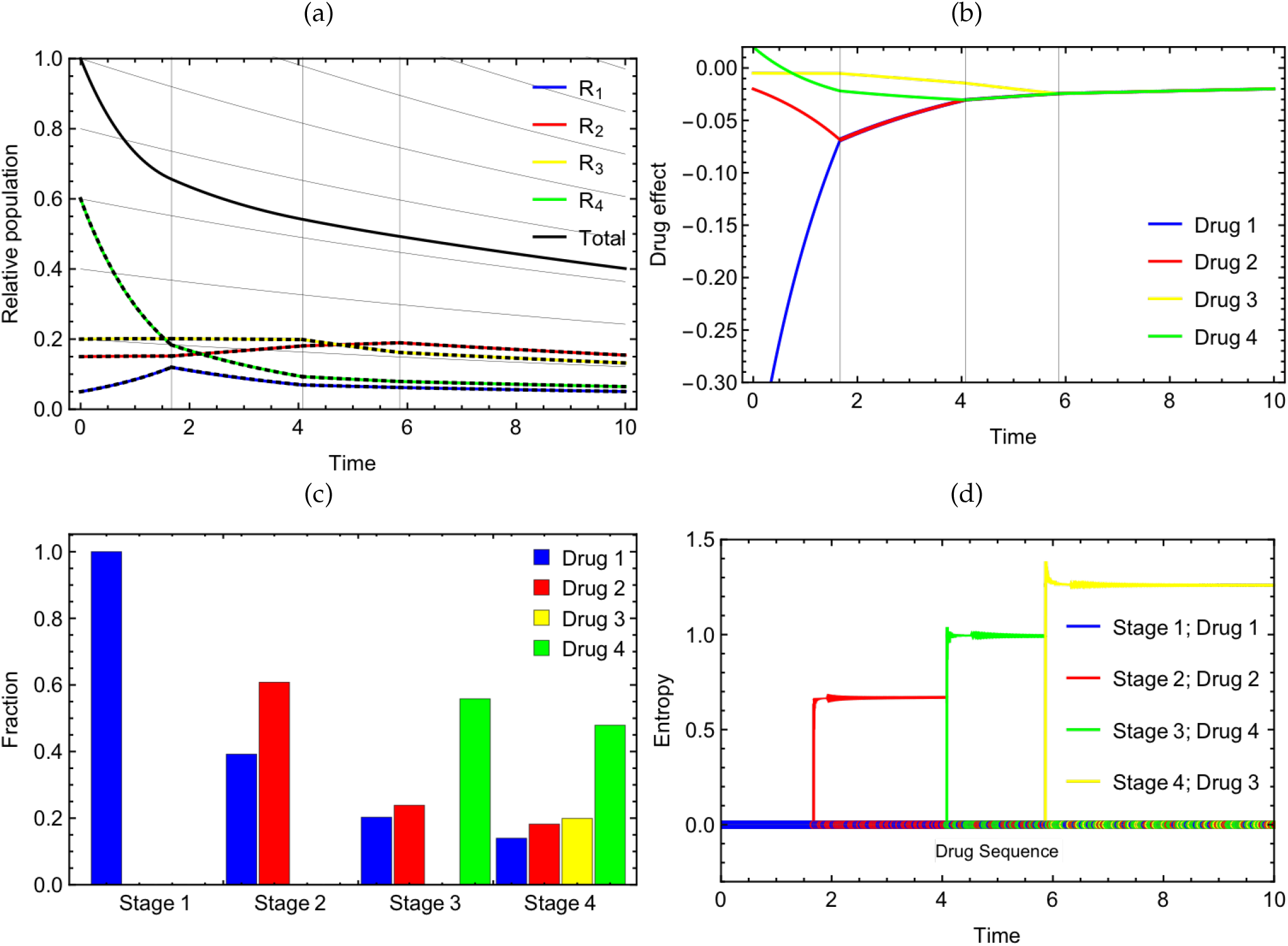
Optimal therapeutic dynamics with a cycle of four asymmetric drugs. Same types of plots with Figure 5 with four asymmetric drugs: 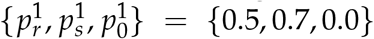, 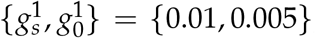, 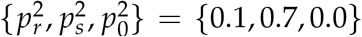, 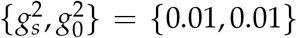, 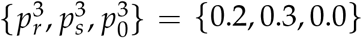, 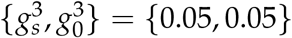, 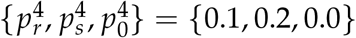, 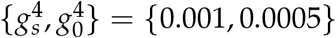, 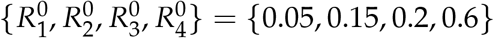

In both cases simulated by the algorithm (Figure 4 (a)), there are up to *N* “stages" of time periods in the optimal administration. (Figure 5, 6 (a)) We define a stage as a time interval in which a same drug combination is involved. Starting initially with the most effective drug(s) of the first stage, with progression to subsequent stages, one or more additional effective drug(s) is chosen to start, and eventually at the last stage all the drugs are activated. (Figure 5, 6 (b,c)) The choice of the additional drugs and duration of the stages except the last stage depend on the initial proportions of subpopulations and model parameters. This qualitative change transitioning to the last stage is because by this time, the heterogeneity has been effectively shaped by the previous stages of drug administration. The last stage starts with a specific subpopulation makeup 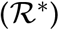 that can be represented by the following expression, if 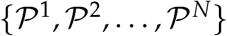 is linearly independent. (See by Theorem A.7 in Appendix.)

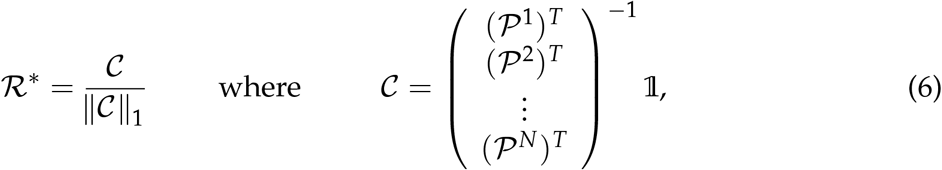

or simply,

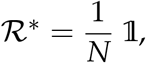

in symmetric drugs (Corollary A.8). There exist possible parameter values which do not hold the linear independence property, if they satisfy one of several strict conditions like Theorem A.9. However, our analysis will focus on other general cases that the expression (6) can be utilized.

The last stage lasts as long as therapy is necessary, keeping the same cellular makeup meanwhile.

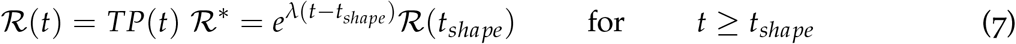

All the subpopulations and total population of the last stage are presumed to change exponentially (See the comparison between the population curves and the light-gray guideline curves on Figure 5, 6 (a).). For symmetric cases, it has been proved in Theorem A.10, that the exponential rate is the average of proliferation rates of all subpopulations,

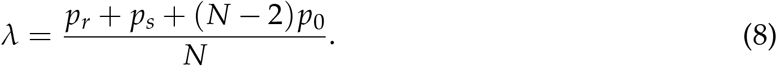

All stages except the last one choose better drugs and gradually level the effectiveness of all drugs evenly, so we will call those finite stages the “shaping period". On the other hand, in the last stage, drugs are equally effective for the shaped heterogeneous tumor, with all drugs being given to continue this balanced effect (adaptive therapy period). The finding of the optimal therapeutic regimen is consistent with our previous work [1] of 2 drugs, in that (i) at each time point the best strategy of drug administration requires the drug(s) with the highest current effectiveness, and (ii) the optimal administrations consist of a shaping period and a period of adaptive therapy.

In a symmetric drug regimen, after the shaping period, all the subpopulations maintain equal density (Stage 4 of Figure 5 (a)) with same intensity of drugs (Stage 4 of Figure 5 (c)) given in turn. However, in the other stages in the case of symmetric drugs, and all stages in the case of asymmetric drugs, subpopulation makeup, relative drug intensities, and the duration of the stages are nonuniform and infeasible to derive analytically. We can clarify them only through numerical methods given drug parameters by running the System (5) in a discrete way as we did for the examples in Figure 5, 6. For reference, in our previous work [1], such quantities were able to clarify explicitly, since it only concerned two drugs.

Additionally, we have measured the Shannon entropy of the drug indices in the sequence of drugs chosen along the algorithm in our examples. Despite the aperiodicities we observed in the sequences, the entropy along the optimal trajectories are more or less consistent, i.e., flat entropy curves in each stage; Figures 5, 6 (d)) within each stage. This allows us to infer that the sequences are almost periodic, and drug prescription does not significantly differ within these stages. Inspired by this observation, we plotted the dynamics of instantaneous drug cycles (Equation (2)) using the relative drug assignments of each stage found at Figures 5, 6 (c). The relevant curves are indicated by the dashed curves in Figures 5, 6 (a). Based on their consistency to the algorithmic histories on the same panel, we can presume that the drug cycle would be periodic, but errors generated by the practical nonzero time step 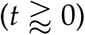 result in minor aperiodicities.

## 4 Practical guidelines for optimal drug administration

Though we have derived the optimal therapeutic regimen for symmetric and asymmetric drug cycles in the previous section, there are practical barriers to applying this regimen in a clinical setting. It is unlikely that the drug parameters and initial tumor status as defined in our algorithm are known for real drugs and a real tumor in a real patient. Therefore, we propose a method of treatment to approximate this optimal therapy in two cases: when the sub-populations of each cell type are known and when only the total tumor cell population is known.

In each case, we use one or more ‘testing periods’ each lasting *N*Δ*t* where Δ*t* is the smallest possible period of single drug administration, and in reality, response measurement. In the *N* successive Δ*t*-long time windows, all of the *N* drugs are given in turn, and the available population data is measured at the end of each window. After this procedure is performed, we will have discrete population data at *N* + 1 time points

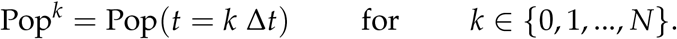

We base our ‘realistic’ strategy around total tumor population because of the recent explosion of robust techniques to obtain this information in a clinically relevant setting [22]. In this particular method plasma cell-free DNA is sampled from a patient with relatively high temporal frequency and used to resolve the corresponding evolutionary dynamics. We use these techniques as a benchmark for clinically leverage-able tumor population data which can be implemented in an algorithm as we describe below.

### 4.1 Available data: subpopulations 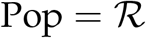

Implementing the subpopulation data over just one single testing period (explained above), we derive several equations which represent the relationships between the data and drug parameters, and then, to find the parameter values. We will approach those problems for two conditions separately: two drugs *N* = 2, and three or more drugs *N* ≥ 3.

In the case of two drugs, there are only two subpopulations 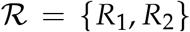, and three parameters for each drug *i* ∈ {1,2} which are two proliferation rates 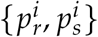 and one transition rate from sensitive to resistant types *g^i^*. No subpopulation or parameters related to neutral cells is included in the system of two drugs. Solving the differential system (1) with the initial conditions, we have the following equations for each Δ*t*-long time window.

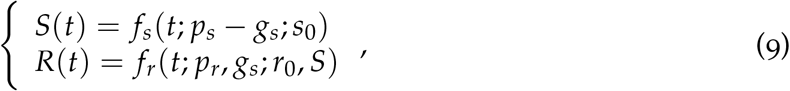

where 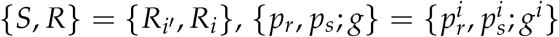 and 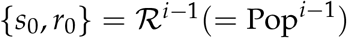. Applying the data of next time point 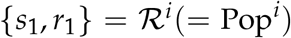 to (9), we can specify the equations with drug parameters and known values only.

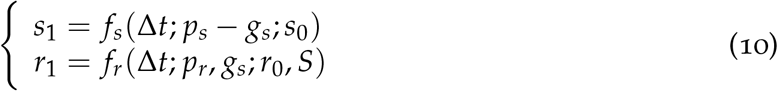

Similarly, in the case of a larger number of drugs *N* ≥ 3, we can solve (1),

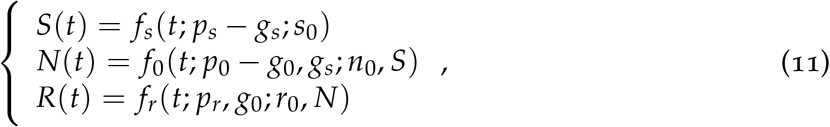

where 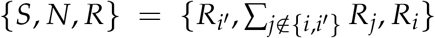 and 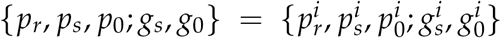. Also, {*s*_0_, *n*_0_, *r*_0_} and {*s*_1_, *n*_1_, *r*_1_} are elements or the summations of elements in *Pop*^*i*−1^ and *Pop^i^* respectively. Applying {*s*_1_, *n*_1_, *r*_1_} to (11) yields,

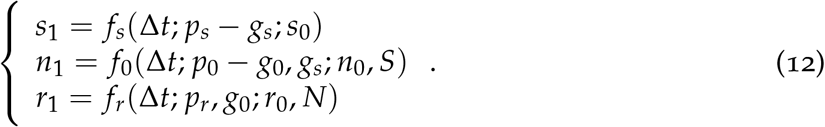

Both (10) for the two-drug cases with three unknown parameters and (12) for *N*-drug cases with five unknown parameters (*N* ≥ 3) are underdetermined. One strategy to resolve this issue in either case is by assuming a specific ratio between proliferation rates and transition rates, i.e., |*p_s_*| = *αg* for 2 drugs and {|*p_s_*|, |*p*_0_|} = *{α_s_g_s_, α*_0_*g*_0_} for more drugs, with a reasonably chosen *α, α_s_, α*_0_ ∈ (0,1) like *α* = 0.1. Then, using the conditions and equations (10 or 12) along with the data from one testing period, we can infer all the drug parameters. The discovered parameters will give the complete information required to run the algorithm of optimal administration (Figure 4 (a)).

### 4.2 Available data: total population only Pop = *TP*

In most cases of cancer, however, detailed information about tumor heterogeneity cannot be detected over time. Clinically, total tumor size is the highest resolution data that can be reasonably (and even then poorly) measured. For such cases with limited data, rather than trying to solve the differential equations as shown in the previous section, we tried to approximate drug effects using the levels of changes of the total population data of testing period, and checked which drug(s) are most reasonable to prescribe. The chosen drug or drugs is continually prescribed as long as it is effective, and its effects are continuously monitored. When the chosen drug loses efficacy, we need to select drugs again. Since we will not know sub-populations and drug parameters, the equation (4) is not applicable. Instead, another (*N*Δ*t*)-long testing period is required to choose the drugs of the next ‘round’. Therefore, in our algorithm of optimal prescription only with total population monitored, we suggest to pair up “testing" period and “therapy" period, and repeat them (See Figure 7 (b)). Even though the testing period can transiently worsen outcomes compared to standard therapies for a finite period of time, it is the only way to ascertain the relative strength of each drugs selection pressure. So, we strived to reduce the time taken for each testing period. One way we implemented this in our algorithm was to choose and apply multiple effective drugs, rather than one single best drug. We chose this strategy because utilizing a strict criterion of drug selection would not allow for currently chosen drugs to be effective for a long time and would require a change of drug more frequently.

**Figure 7:**
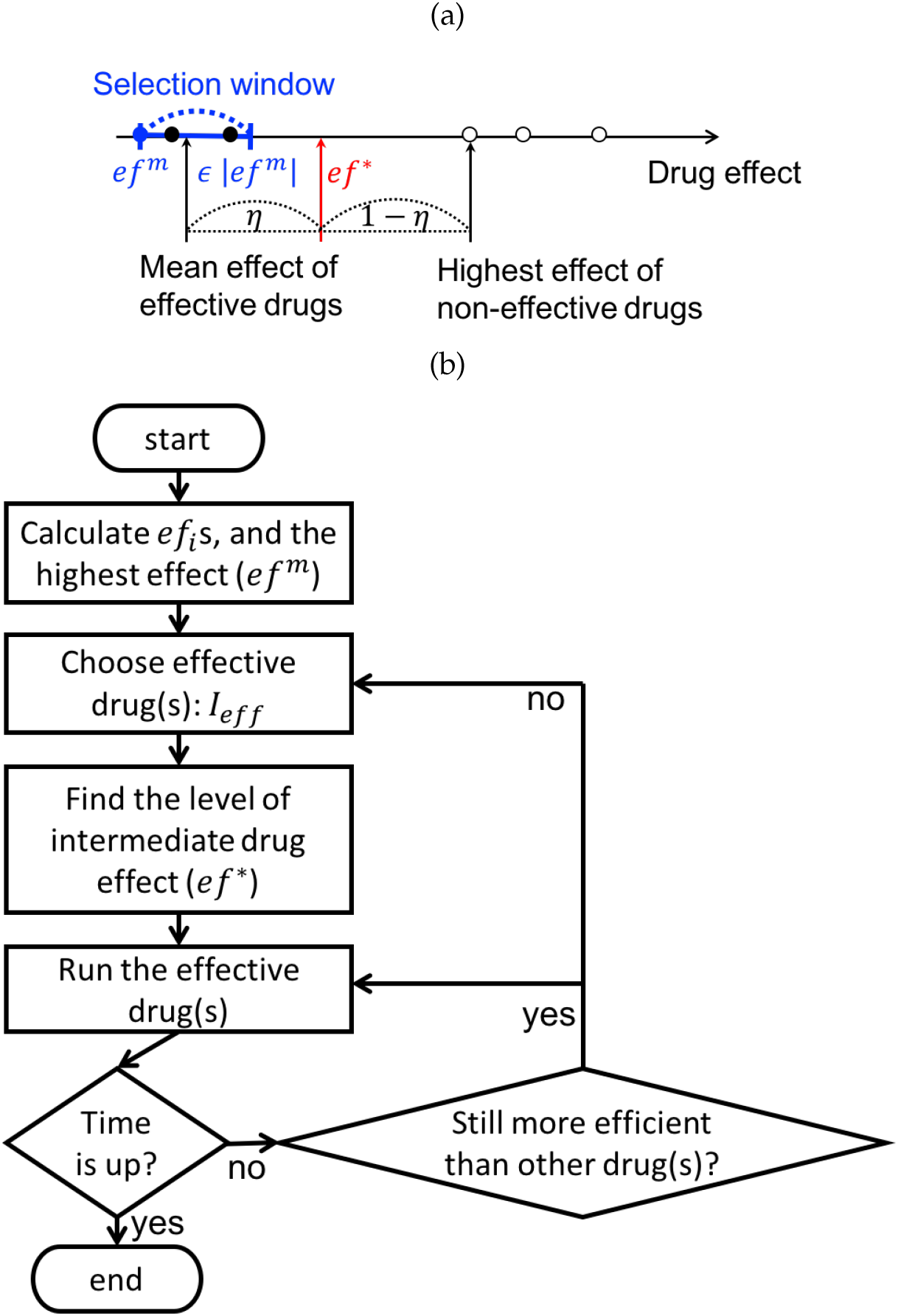
Algorithm for the administration of the optimal drug schedules when total population data is available, but not subpopulation nor drug parameters. (a) Diagram of the classification of “effective" and “ineffective" drugs, and the threshold of drug effectiveness (*ef^*^*). The effectiveness levels of the most effective drug (*ef^m^*), effective drugs and ineffective drugs are indicated by blue-filled circle, black-filled circles and empty circles, respectively. (b) Flow chart of the optimal therapy algorithm based on the “effective" drugs from (a).

Now, let us describe our algorithm for optimal therapeutic prescription (Figure 7 (b)) with only total tumor cell population trackable. We measure the effect of *Drug i* (*i* ∈ {1,2,…, *N*}) by the approximate derivative,

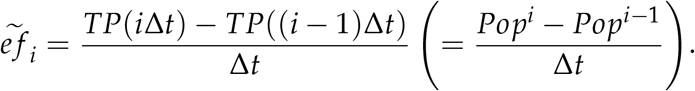

From the evaluated 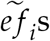, we choose the effective drug(s), based on the current status of the tumor,

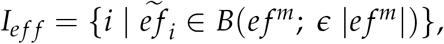

based on the highest effect,

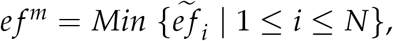

and the parameter of effective drug interval (*ϵ*). Another important quantity included in the algorithm is the threshold of the drug effectiveness necessary to determine if we can keep using the current drug(s) or need to re-select drug(s),

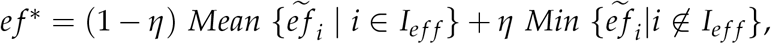

which is between the average effect of “effective" drugs, and the highest effect of “ineffective" drugs. In our study, we simply fixed the threshold parameter, *η* = 0.5 (See Figure 7 (a) for the visualized explanations about the values and parameters in the algorithm.).

Our algorithm of approximated optimal therapy was compared to the actual optimal therapy by measuring the area between population curves, in two ways. Using total population curves, we measured

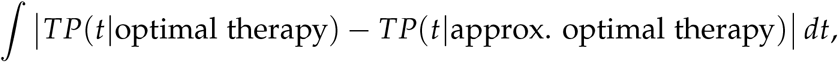

and using subpopulations curves,

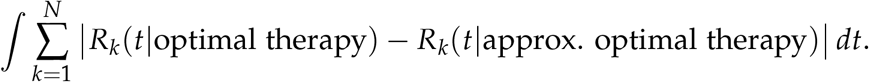

Using both methods, we found that both strategies which were too strict or too generous do not generate tumor histories close enough to the optimal one (See Figure 8 (b)). A proper level of drug selection window is necessary, like our example case of Figure 8 in which the approximation is reasonably close with *ϵ* ≈ 0.03 (See Figure 8 (a)). It is an expected observation, because a strict drug selection strategy will require unnecessarily frequent testing periods, and a generous strategy will include barely effective drugs as well in the treatment.

**Figure 8:**
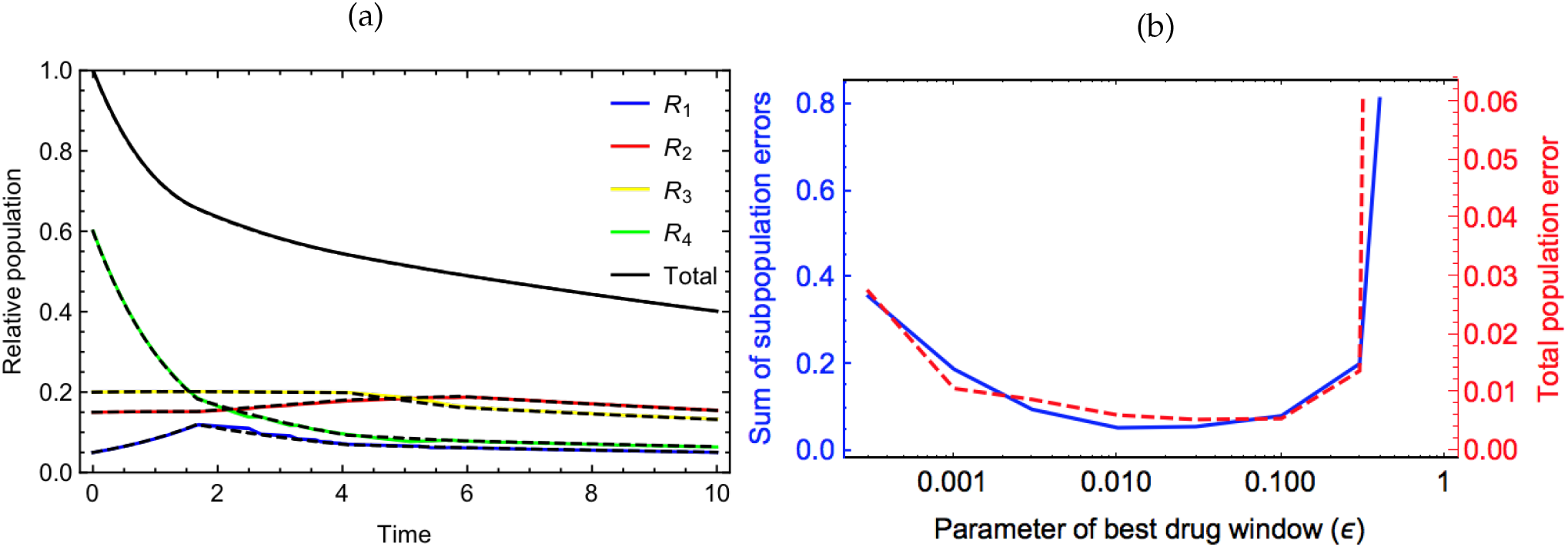
Comparison between the realistic approximation of the optimal therapy (by the algorithm on Figure 7 (b)), and the actual optimal strategy. (a) Optimal therapeutic effect generated by the practical algorithm assuming that only total population is trackable (solid curves) with *ϵ* = 0.03 and *η* = 0.5, compared to the effect of the optimal therapy in theory (dashed curves). Other used parameters and initial tumor status are the same as in the example in Figure 6. (b) Errors between the approximated and actual optimal histories, over the range of the parameter of effective drug window.

## 5 Conclusions and discussion

The phenomena of collateral sensitivity presents an opportunity to improve effectiveness in drug therapy without the need for drug discovery, a process that requires enormous amounts of time and money. Taking advantage of CS clinically however, requires a better understanding, of the evolutionary dynamics under changing therapy. To address this, we developed a mathematical model of ordinary differential equations describing the effect of a collaterally sensitive drug cycles, and explored the optimal treatment regimens within the confines of this simplified model. Consequently, we found that the optimal therapeutic effect can be derived when we switch drug to the best-effect-drug at every moment. While this is somewhat intuitive, choosing the timing, and order of switching is a difficult prospect given the lack of perfect data in clinical settings, and given the heterogeneity which is hallmark of cancer.

In our ODE model, drug switching is implemented by changing drug-dependent parameters 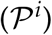 and transitions between cell types. In accordance with the ‘optimal’ drug schedule, drugs are switched instantaneously after the first stage of single-drug therapy. However, such rapid switching is not feasible clinically. To address this, we developed a time-series algorithm mimicking the ODE system and the nearly instantaneous switches (Figure 4 (a)). As expected, the algorithm-based simulation is smooth with a small time step (Figure 4 (b)) and shows a good consistency with the ODE system (Figure 5, 6 (a)).

The order of the drug sequence in each stage of our example simulations is not exactly periodic, even though the pattern of periodicity is (at least vaguely) observed (Figure 5, 6 (d)). Also, in the last stages where all the drugs are involved, *Drug i* is not always followed by *Drug i′*. However, the drug combinations of a complete drug cycle is needed to control tumors. Specifically, if there is a flaw in suppressing one type of cell population, even if its population was very small at the beginning, it can exponentially increase at the end (Figure 9) – much like the concept of evolutionary escape.

**Figure 9:**
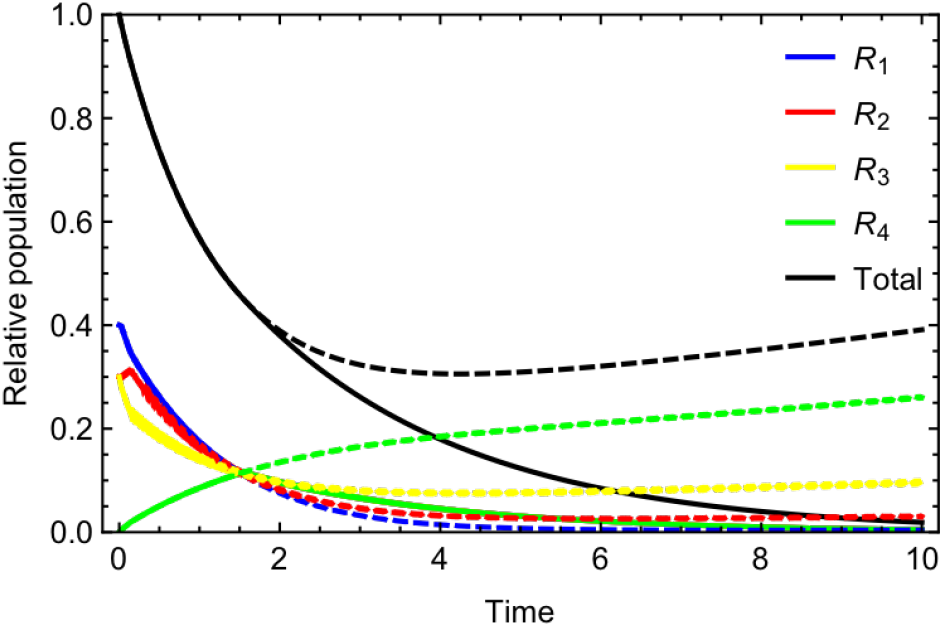
Comparison between the effects of a complete collaterally sensitive drug cycle and an incomplete cycle. The complete cycle is comprised by all the four drugs (relevant to the solid curves), and the incomplete cycle is by only *Drug* 2, *Drug* 3 and *Drug* ¦ with *Drug* 1 being dropped (relevant to the dashed curves). Used values: {*p_r_, p_s_, p*_0_} = {1.0,2.5,0.0}, {*g_s_,g*_0_} = {0.25,0.25} for all drugs, 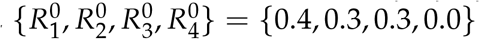

In our research, the measure of drug effect at a specific time point is defined by the instantaneous rate of change in total tumor size under the drug (Equation 4). It is therefore dependent on the heterogeneity in the tumor comprising the tumor at a given moment and how effective the drug is to each cell type. Hence, if the cell composition and/or drug parameters are unknown, we cannot measure the drug effect and therefore the optimal drug switch timing cannot be captured. For such cases, which is the majority of cancers, we developed an algorithm within which drugs are selected based on only total population (Figure 7(b)). Cell population dynamics with this algorithm show a good consistency with the fully informed optimal therapeutic strategy (Figure 8 (a)). The traditional time gaps required to obtain updated diagnostics and prescription is several weeks or months, [22] which would be too long to expect a result close to an optimal solution as we aim for here. Based on the results of our algorithm simulation (Figure 8 (a)), it is evident that performance of the algorithm degrades with increasing time between updates – suggesting an avenue of study to formally optimize the costs and benefits of this time, though that is beyond the scope of this work.

There are many collateral sensitivity relationships found among antibiotics [27, 34] and anticancer [13] drugs. A collaterally sensitive drug cycle is a chain of such relationships with various lengths (> 2) [13] or more. It is possible that the increased sensitivity shown after another drug could be a temporal phenomenon happening in a specific status, which is too complicated to recapitulate within this structure. Also, even if a part of the cell population follows the dynamics of a rotating resistance and sensitivity pattern, there could be many more types of cells not involved in any cell types within this structure, which will result in treatment failure, as we discussed in Figure 9. A detailed study with antibiotics [27] applied empirical fitness measurements to assign cellular classifications with genes and drug-induced transitions among the genotypes. Finding best-case therapy ordering in multi-drug scenarios was also studied by Maltas et al. without reliance on the underlying fitness landscape, somwhat akin to our “imperfect" clinical information, and found that therapy could be improved. [35]

Each of these previous studies assume that the dynamics measured at the beginning of the study do not change through time. Given the importance of (epi-)mutation, and not just changing frequencies, it is uncertain on what timescales this is a fair assumption. [9] Another simplification we have made is to assume that there is no interaction between cell-types, something which has been shown in at least breast [36], lung [17] and pancreatic cancers [37]. However, we believe our cell classification structure is meaningful in that it reflects collateral sensitivity phenomena reasonably and the simplicity of the corresponding linear ODE system enables analytic study.

In addition, our system and corresponding a analysis lend well to the theoretical and clinical framework of adaptive therapy [2]. In adaptive therapy, drug administration is alternated with drug holidays to minimize tumor growth and resistant cell population. The central issues of optimal time for drug switching to maintain an optimal sub-population structure is apropos to the issues explored in this and our own previous work. Adaptive therapy has demonstrated clinical utility in markedly improving tumor response for a subset of cancers in theory, [3] in pre-clinical models [38] and clinically [39]. While this purely theoretical study should not be used to inform clinical care at this time, [40] we submit that further application of the techniques herein present a way forward for precise timing of drug switching to further optimize single drug adaptive therapy and higher order drug sequences alike.

## Appendix A Differential system of instantaneous drug switch

**Definition** In addition to the definitions from Table 1, we defines further notations to facilitate the descriptions on proofs.

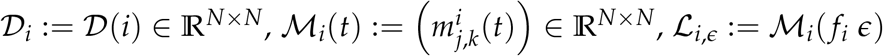

with 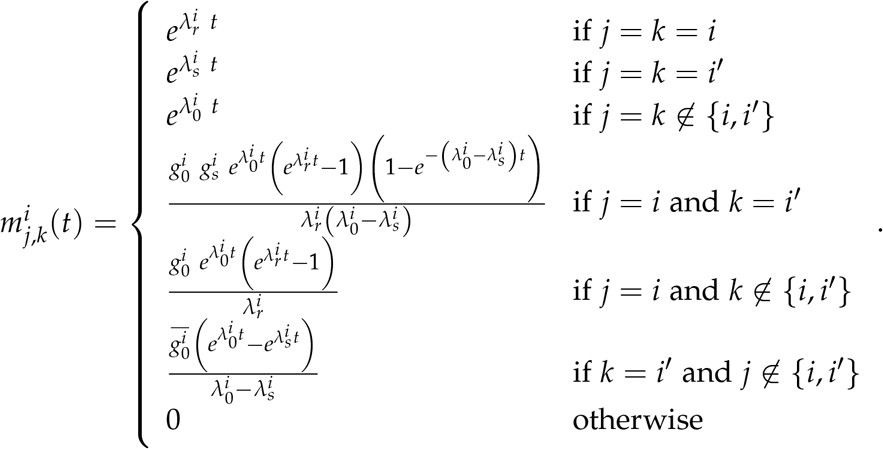

**Proposition A.1.** *Under the therapy with Drug i:*

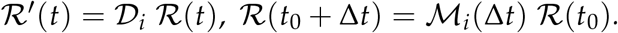

**Proposition A.2.** 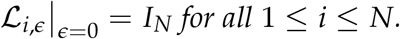

**Proposition A.3.** 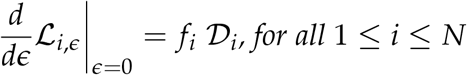

**Lemma A.4.** 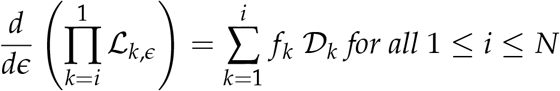

*Proof*.

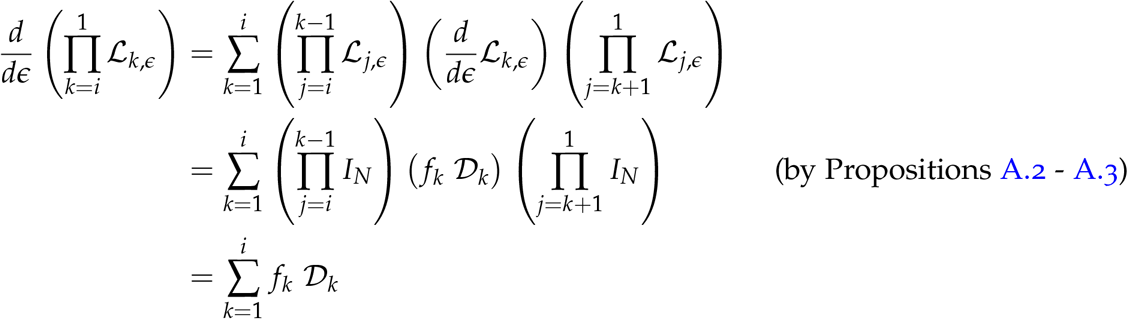

**Lemma A.5.** *For any positive integer, n, and an any integer, i, in* [1, *N*]

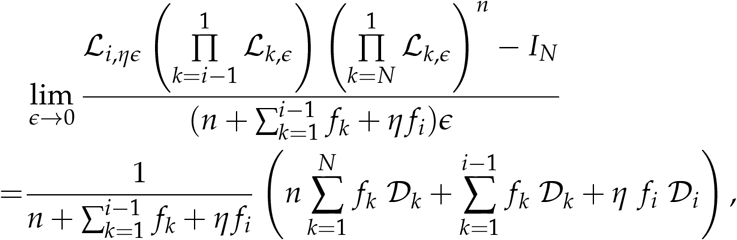

*where η is some number in* [0,1) *and* 0 ≤ *f_k_* ≤ 1 *with* 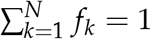.

*Proof*.

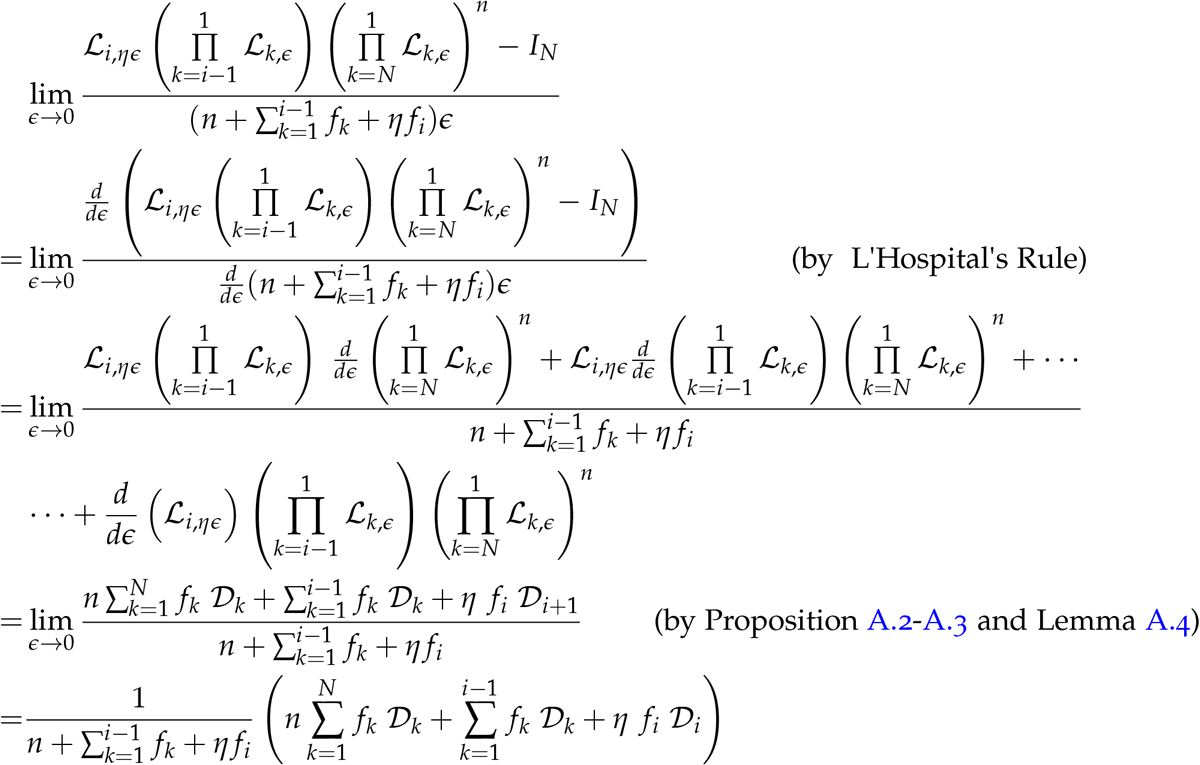

**Theorem A.6.** *If Drug*1, *Drug*2, …, *DrugN, are prescribed in a cycle, and are switched instantaneously with relative duration of* 0 ≤ *f*_1_, *f*_2_, …, *f_N_* ≤ 1 (*where* 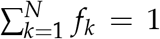*), respectively*, 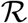 *obeys*

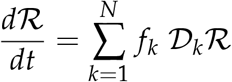

*Proof*. For any time point *t*_0_, let us define 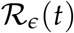 as a vector-valued function of *R*_1_ (*t*), *R*_2_(*t*),…, and *R_N_*(*t*) describing the cell population dynamics under a periodic therapy starting at *t*_0_ with *Drug i* assigned for 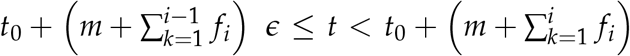 where *m* is any non-negative integer and *i* is any integer in [1, *N*].

For any Δ*t* > 0, there uniquely exist *n* ∈ {0,1,2,…} and *η* ∈ [0,1) such that 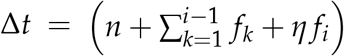. Then, by Proposition A.i and the definitions of 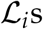,

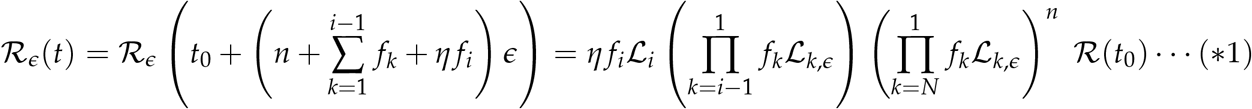

where 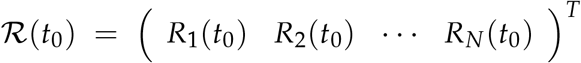. And, 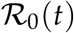 represents instantaneous drug switching.

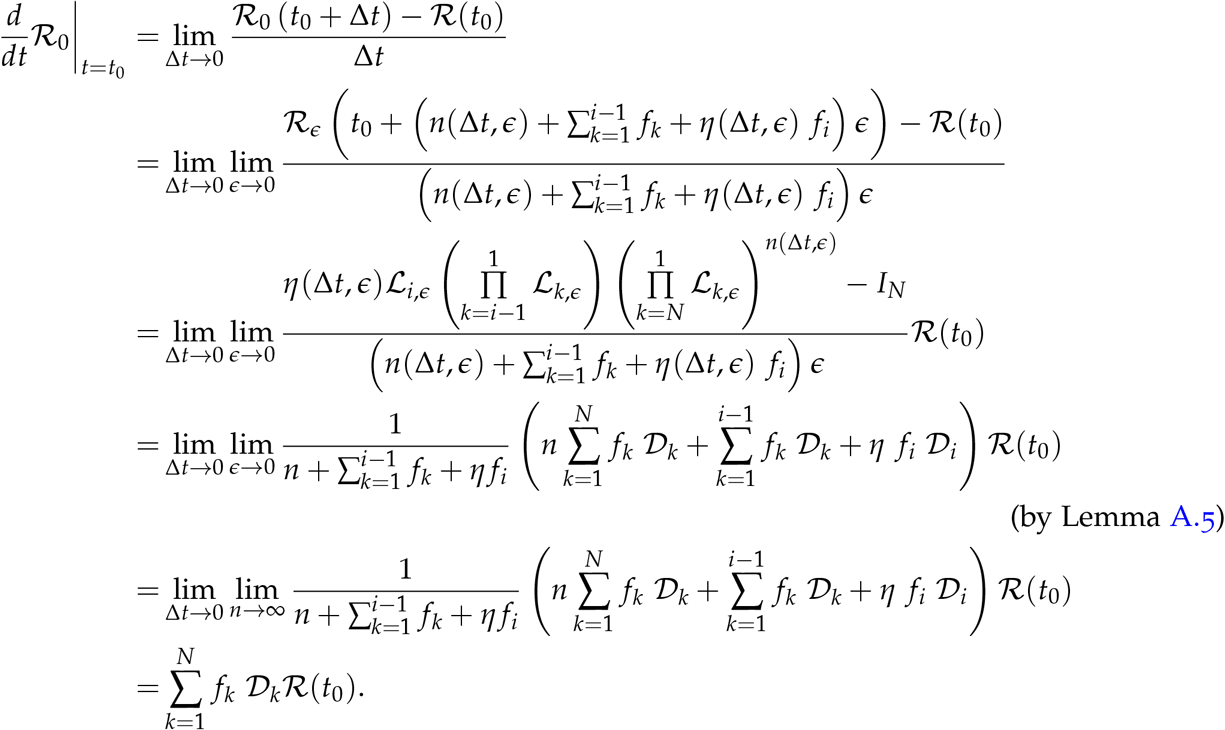

Therefore,

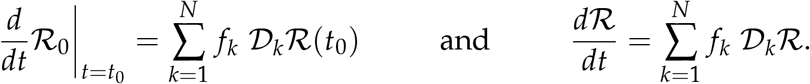

**Theorem A.7.** *The population makeup at which all the drugs are equally effective is*

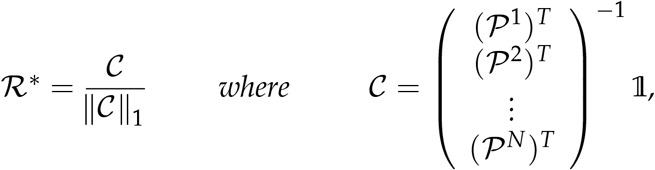

*where* 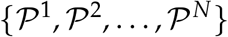 *is linearly independent*.

*Proof*. By the definition (4), the effect of *Drug i* is

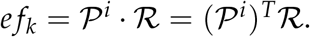

Then, at a specific population makeup with balanced drug effects, denoted by 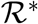,

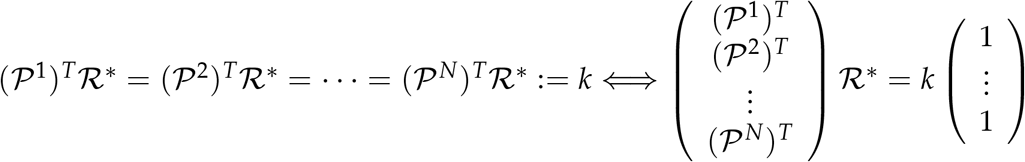

where *k* is a constant representing the level of balanced drug effect. Since 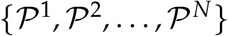 is linearly independent, we can isolate 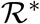 with an inverse matrix.

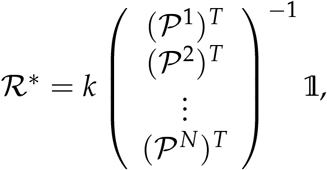

Also, 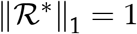, therefore

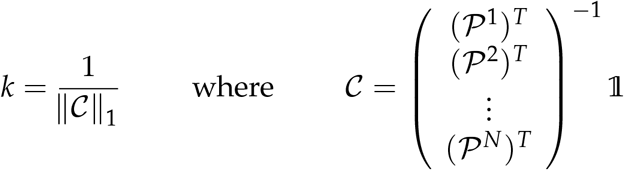

and

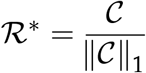

**Corollary A.8.** *When the drugs are symmetric, the population makeup at which all the drugs are equally effective is*

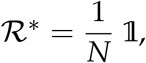

*where* 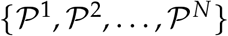 *is linearly independent*.

*Proof*. Similar to the proof of Theorem A.7, there exists unique solution of 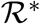 and *k* satisfying

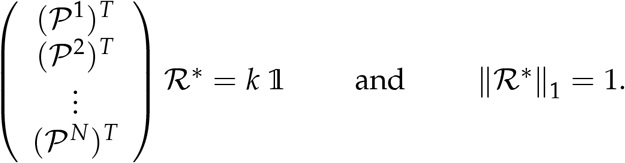

If we plug in 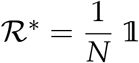 to the equation,

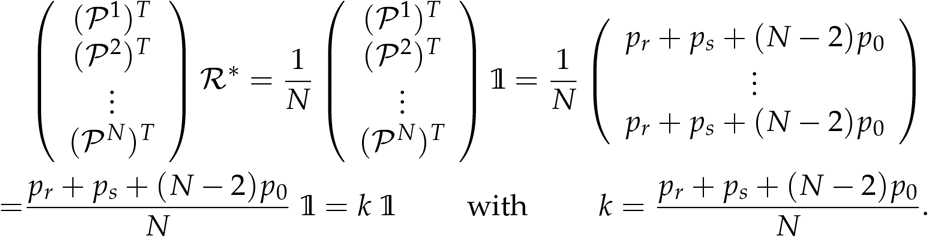

Therefore, proved.

**Theorem A.9.** 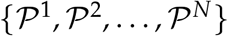 *is linearly dependent, if the drugs are symmetric (i.e.*, 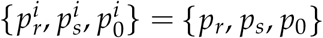 *for all i) and p_r_ + p_s_* + (*N* − 2)*p*_0_ = 0

*Proof*. It suffices to show that there exists a linear combination of 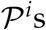 that equals to a zero vector. Let the general form of a linear combination be 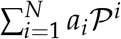, where *a_i_*s are constants. If *a_i_* = 1 for all *i*, 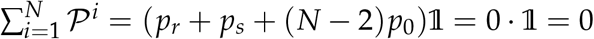. Therefore, 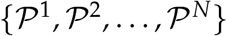 is linearly dependent.

**Theorem A.10.** *Under instantaneously switching of symmetric drug cycle, total population with equal drug duration* (*f*_1_ = *f*_2_ = … = *f_N_* = 1/*N*), *TP, is changing exponentially with the growth/decay rate*,

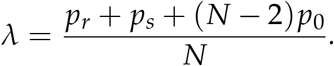

*Proof*. The derivative of total population, *TPD*, is the summation of the vector, 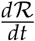

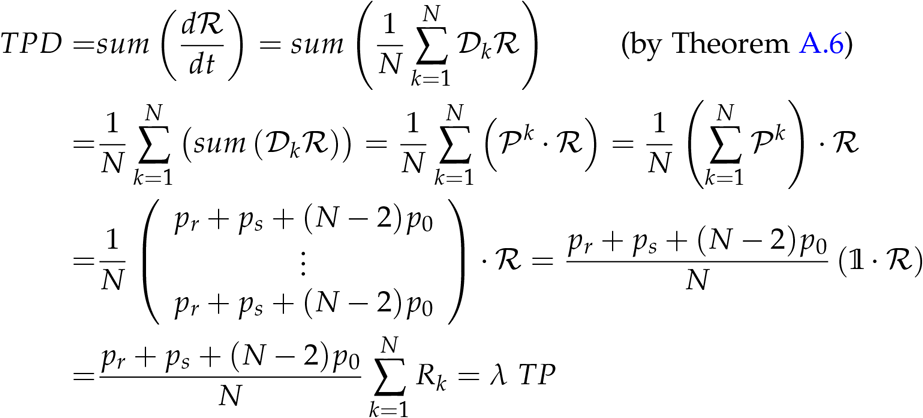

## Notes

### Competing Interest Statement

The authors have declared no competing interest.

### Summary of Updates

Title changed

